# Pitfalls in SARS-CoV-2 PCR diagnostics

**DOI:** 10.1101/2020.06.03.132357

**Authors:** Kerstin Wernike, Markus Keller, Franz J. Conraths, Thomas C. Mettenleiter, Martin H. Groschup, Martin Beer

## Abstract

To combat the COVID-19 pandemic, millions of PCR tests are performed worldwide. Any deviation of the diagnostic sensitivity and specificity will reduce the predictive values of the test. Here, we report the occurrence of contaminations of commercial primers/probe sets with the SARS-CoV-2 target sequence of the RT-qPCR as an example for pitfalls during PCR diagnostics affecting diagnostic specificity. In several purchased in-house primers/probe sets, quantification cycle values as low as 17 were measured for negative control samples. However, there were also primers/probe sets that displayed very low-level contaminations, which were detected only during thorough internal validation. Hence, it appears imperative to pre-test each batch of reagents extensively before use in routine diagnosis, to avoid false-positive results and low positive predictive value in low-prevalence situations. As such, contaminations may have happened more widely, COVID-19 diagnostic results should be re-assessed retrospectively to validate the epidemiological basis for control measures.

## Introduction

In December 2019, an outbreak of an unexplained acute respiratory disease of humans was reported in Wuhan, China (WHO, 2020d). As causative agent of the disease now named COVID-19, a novel betacoronavirus referred to as severe acute respiratory syndrome coronavirus 2 (SARS-CoV-2, previously known as 2019-nCoV) was identified (Zhu et al., 2020). COVID-19 rapidly evolved into a global pandemic (WHO, 2020b) resulting in millions of infections and several hundred thousands of deaths. Overall, about 20 % of the symptomatic infections are severe or critical, with much higher rates in the elderly or when certain underlying health conditions exist (WHO, 2020b). However, also asymptomatic infections occur and it is estimated that virus transmission from asymptomatic humans accounts for about half of all COVID-19 cases (He et al., 2020), which might be particularly critical when asymptomatically infected health care workers transmit the virus in hospitals or care homes for the elderly.

Diagnosis is currently based primarily on real-time RT-PCR (RT-qPCR) using nasal or throat swabs. To identify and isolate infected individuals, thereby interrupting transmission chains, millions of RT-qPCR tests are carried out (Hasell et al., 2020). With such a large number of diagnostic tests and yet low prevalences of infected humans, it is of utmost importance to ensure a high level of quality management in the testing laboratories to guarantee an optimal and reliable diagnostic accuracy. Any deviation of the diagnostic specificity of the PCRs, e.g. through contamination of reagents with target sequences, mix-up or cross-contamination of samples will significantly reduce the positive predictive value of the test. Here, we report contamination of commercial primers and probes with oligonucleotides as an example for pitfalls during PCR diagnostics with a drastic effect on diagnostic specificity. This example emphasizes the need for continuous and comprehensive quality management in all diagnostics steps.

## This study

For the detection of SARS-CoV-2 genome, two real-time PCRs listed on the website of the World Health Organization (WHO) (WHO, 2020a) were established and validated in our laboratory. To increase the diagnostic accuracy, systems targeting different genomic regions were selected. The first assay (“E-Sarbeco”) is based on the E gene coding region (Corman et al., 2020) and the second assay (“nCoV_IP4”) targets the RNA-dependent RNA polymerase (RdRp) gene (WHO, 2020a). To control for efficient RNA extraction and amplification, both assays were combined with an internal control system based on the housekeeping gene beta-actin (Wernike et al., 2011). Primers and probes were ordered from four different commercial companies in March and April 2020, amongst them major oligonucleotide suppliers on the European market. Both duplex SARS-CoV-2/beta actin real-time PCR systems were validated using two different real-time PCR kits, namely the AgPath-ID™ One-Step RT-PCR kit and the SuperScript III One Step RT-PCR kit (both produced by Thermo Fisher Scientific, Germany) to increase flexibility in case of supply shortage.

As part of our internal quality management, each batch of primers/probe is investigated regarding its sensitivity and specificity using SARS-CoV-2 RNA and negative samples (phosphate buffered saline (PBS) or nuclease-free water) before the oligonucleotides are applied in routine diagnosis. During these pre-tests, the first primers/probe sets from supplier A purchased in March 2020 (set A-1) performed as expected. However, subsequently, very high genome loads were found in some newly purchased E-Sarbeco primers/probe sets. Quantification cycle (Cq) values as low as ~17 or ~22 were measured in negative control samples (table 1) indicating a high level of contamination in reagents obtained from some oligonucleotide suppliers. While the problems in performance are obvious in these cases, there were also primers/probe sets that displayed contaminations only at lower levels. As an example, when we used a separate batch of oligonucleotides from supplier A (set A-2), only two out of 27 negative control samples reacted weakly positive. To exclude the PCR chemistry or the internal control oligonucleotides as potential sources of the false-positive results, samples from the first German proficiency test on COVID-19 diagnostics (INSTAND e. V. and GBD Gesellschaft für Biotechnologische Diagnostik mbH) as well as seven negative RNA isolation controls (RIC = PBS) were tested using the incriminated primers/probes in combination with two different batches of both RT-PCR kits. Every combination, in which the first set of primers/probe was applied, yielded correct results, while the incriminated primers/probe (set A-2) resulted in several false-positive results regardless of the applied PCR chemistry (figure 1). Hence, the primers or the probe were the cause of the false-positive reactions, a phenomenon that seems to occur frequently (table 1). The main reason for the wide distribution of contaminated primers or probes may be the simultaneous production of long oligonucleotides containing SARS-CoV-2 target sequences for real-time RT-PCRs. Especially during the first phase of the establishment and internal validation of SARS-CoV-2 specific real-time RT-PCRs, such oligonucleotides have been widely used as positive controls and were produced by many primer/probe suppliers (Mögling et al., 2020).

**Table 1:**
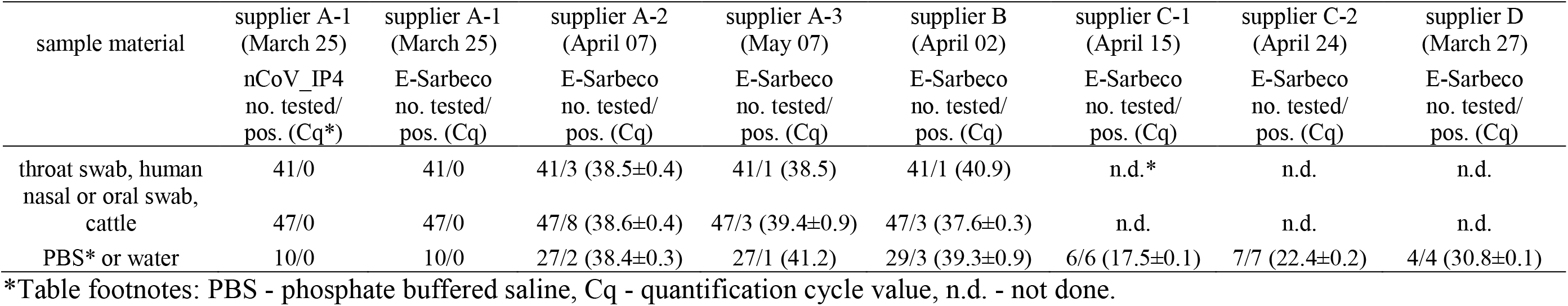
RNA preparations from SARS-CoV-2 negative human throat swabs, bovine nasal or oral swabs and further negative controls (phosphate buffered saline (PBS) or nuclease-free water) were tested by different batches of the identical in-house primers and probe (Corman et al., 2020). The primers/probe sets are named according to the company at which they were synthetized, the delivery dates are given in brackets. When several sets were ordered at the same supplier, they are consecutively numbered. The mean quantification cycle values (Cq) including standard deviations for the false positive results are given in brackets.

| sample material | supplier A-1<br>(March 25)<br>nCoV_IP4<br>no. tested/<br>pos. (Cq*) | supplier A-1<br>(March 25)<br>E-Sarbeco<br>no. tested/<br>pos. (Cq) | supplier A-2<br>(April 07)<br>E-Sarbeco<br>no. tested/<br>pos. (Cq) | supplier A-3<br>(May 07)<br>E-Sarbeco<br>no. tested/<br>pos. (Cq) | supplier B<br>(April 02)<br>E-Sarbeco<br>no. tested/<br>pos. (Cq) | supplier C-1<br>(April 15)<br>E-Sarbeco<br>no. tested/<br>pos. (Cq) | supplier C-2<br>(April 24)<br>E-Sarbeco<br>no. tested/<br>pos. (Cq) | supplier D<br>(March 27)<br>E-Sarbeco<br>no. tested/<br>pos. (Cq) |
| --- | --- | --- | --- | --- | --- | --- | --- | --- |
| throat swab, human | 41/0 | 41/0 | 41/3 (38.5±0.4) | 41/1 (38.5) | 41/1 (40.9) | n.d.* | n.d. | n.d. |
| nasal or oral swab,<br>cattle | 47/0 | 47/0 | 47/8 (38.6±0.4) | 47/3 (39.4±0.9) | 47/3 (37.6±0.3) | n.d. | n.d. | n.d. |
| PBS* or water | 10/0 | 10/0 | 27/2 (38.4±0.3) | 27/1 (41.2) | 29/3 (39.3±0.9) | 6/6 (17.5±0.1) | 7/7 (22.4±0.2) | 4/4 (30.8±0.1) |

\*Table footnotes: PBS - phosphate buffered saline, Cq - quantification cycle value, n.d. - not done.

**Figure 1:**
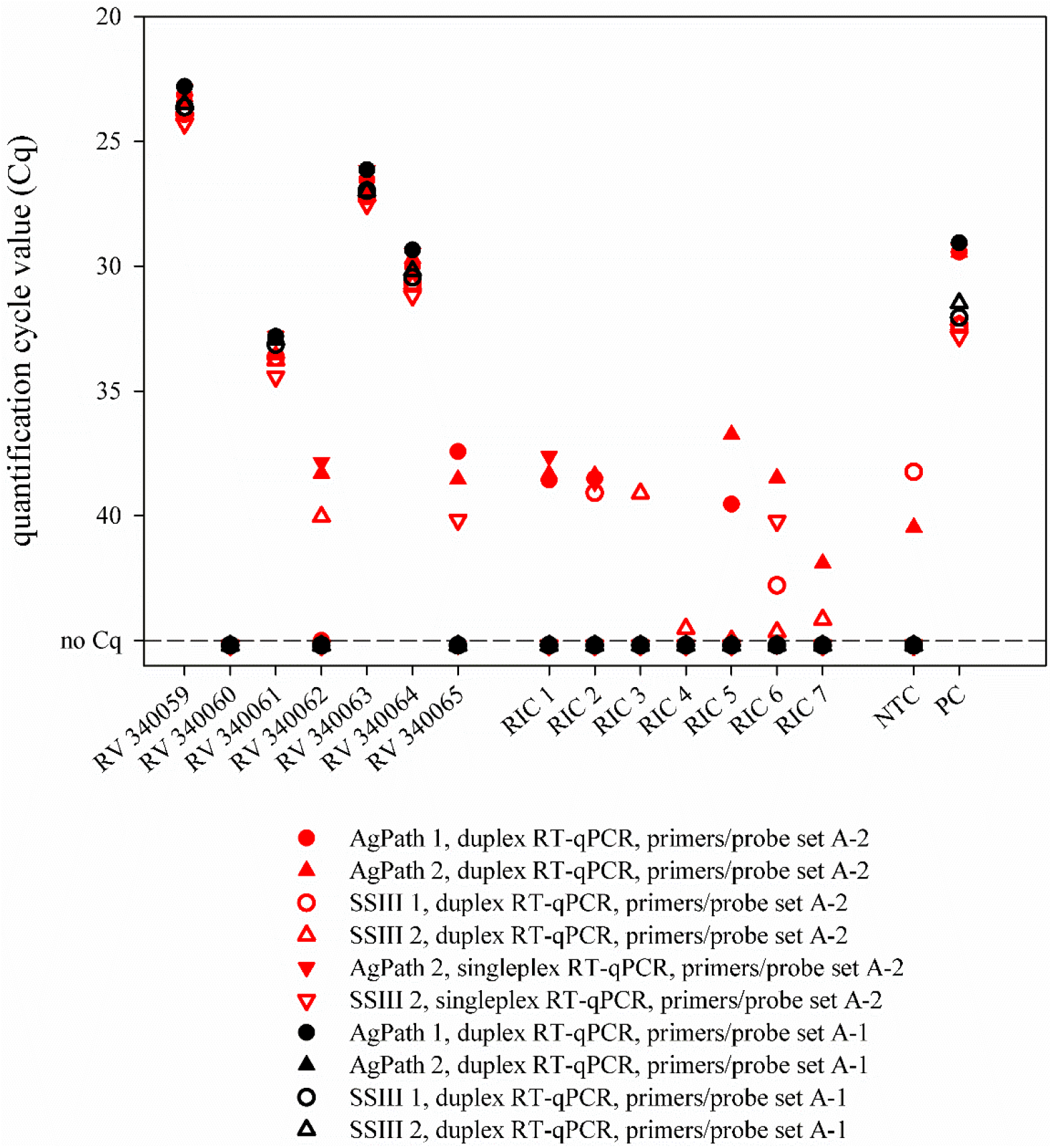
Real-time RT-PCR results generated by using two different batches of the identical in-house primers and probe (Corman et al., 2020) in combination with two distinct PCR kits. RIC - RNA isolation control, NTC - no template control, PC - positive control, AgPath - AgPath-ID™ One-Step RT-PCR kit (Thermo Fisher Scientific, Germany), SSIII - SuperScript III One Step RT-PCR kit (Thermo Fisher Scientific, Germany)

To investigate the impact of the low-level primer/probe contamination on the diagnostic specificity, 41 human throat swabs were tested with the different primers/probe sets A-1, A-2, A-3, and B. Swabs were collected in 1 ml PBS and total nucleic acid extracted from this swab medium either manually (QIAamp Viral RNA Mini, Qiagen, Germany; extraction volume 140 μl) or automated (NucleoMag VET kit, MACHERY-NAGEL GmbH & Co. KG, Germany; extraction volume 100 μl). To exclude nonspecific reactions, which could be caused by other human coronaviruses potentially present in the throat swab samples, 47 oral or nasal swabs of bovine origin (taken before the SARS-CoV-2 pandemic) were included. These specimens represented routine submissions to the Friedrich-Loeffler-Institut, Federal Research Institute for Animal Health, or originated from an unrelated animal trial (Wernike et al., 2018). Positive predictive values were calculated using EpiTools (https://epitools.ausvet.com.au/predictivevalues).

All human and bovine swab samples scored negative by the nCoV_IP4 assay and the first E-Sarbeco primers/probe set delivered at the 25^th^ of March 2020 (set A-1) (table 1). However, when tested by the oligonucleotides A-2, A-3 and B, a total of 13, five and seven of the negative samples scored positive, respectively. Since the empty control (NTC = nuclease free water), which was included in the PCR runs, reacted negatively as expected, the PCRs would have been considered valid during routine diagnostics. Thus, the samples would have been incorrectly diagnosed as positive in settings, where no cut-off for positivity is defined.

If we assume a best-case scenario for specificity based on these results for the A-3 or B / E-Sarbeco setting, the diagnostic specificity was calculated as 0.9756 (40/41; table 1). In calendar week 14 of 2020, 36,885 out of 408,348 samples (9.0%) tested positive in Germany (Robert-Koch-Institut, 2020). Under these conditions, the positive predictive value of the test system was 0.802, i.e. almost 20% of the positive results would have been false-positive. In calendar week 19, 10,187 out of 382,154 samples (2.7%) tested positive. In this scenario, a test system with a diagnostic specificity of 0.9756 had resulted in a positive predictive value of 0.5319, i.e. almost half of the positive results would have been false-positive. Obviously, any further reduction of the prevalence of SARS-CoV-2 infections will result in decrease of the positive predictive value if the specificity of the employed assays is not dramatically increased.

Not only in-house PCRs need to be thoroughly validated in every laboratory, but also commercial kits (Rahman et al., 2020), as they may contain similar primer/probe mixes and produce incorrectly positive results, which will also result in a low positive predictive value.

As an additional component of quality assurance, the preparation of small sample pools might be considered in areas or scenarios with low prevalences (e.g. among asymptomatic persons), which conserves resources and increases sample throughput (Abdalhamid et al., 2020, Eis-Hübinger et al., 2020, Yelin et al., 2020). Most importantly, when such a pool scores positive, all samples need to be re-tested individually, where at least one individual sample should result in the same or a higher virus load than the sample pool itself. In the case of contaminations as described above, the pool will show implausible results during follow-up testing markedly increasing the diagnostic specificity. The WHO recommends widespread testing to combat the COVID-19 pandemic (WHO, 2020b). However, the capacity of SARS-CoV-2 for explosive spread has not only overwhelmed weaker health systems, but also challenges diagnostic capacities (Hasell et al., 2020, WHO, 2020b). Where testing capacity cannot meet the needs, even a prioritization of testing has to be implemented (European Commission, 2020, WHO, 2020c). In such settings of limited resources, pooling of samples might be an option for the serial screening of e.g. asymptomatic health care workers, which is highly recommended to prevent nosocomial transmission of the virus (Rivett et al., 2020). Here, the samples of the German proficiency test (INSTAND e. V. and GBD Gesellschaft für Biotechnologische Diagnostik mbH) were tested in pools consisting of the respective ring trial sample and four negative human throat swabs. The values obtained from the pools were about 2.2 Cq higher than the values of the respective individual samples, but the final assessment was always correct, i.e. each positive sample was correctly identified (figure 2). While there is undoubtedly a (minor) decrease in analytical sensitivity, the pooling option needs to be carefully considered in the light of the current epidemiological situation, as every positive pool needs to be dissolved anyway to test the samples individually. Nevertheless, to screen certain groups, in which the expected prevalence of positive samples is low, pooling might be a resource- and cost-effective option with a minimal loss of diagnostic sensitivity, but with an increase in diagnostic specificity.

**Figure 2:**
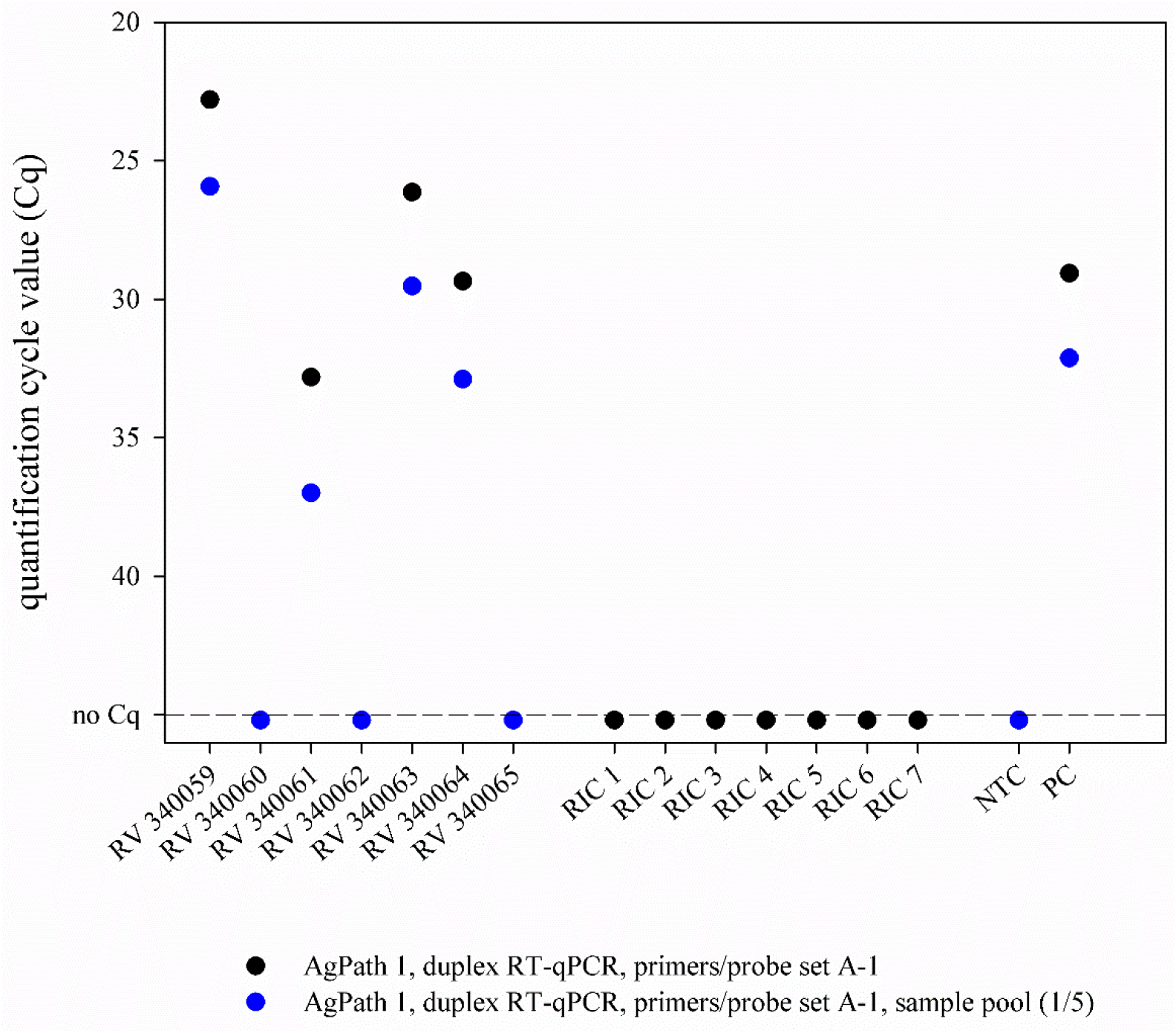
Real-time RT-PCR results of samples that were tested either individually (black dots) or in pools consisting of one SARS-CoV-2 positive and four negative samples (blue dots). RIC - RNA isolation control, NTC - no template control, PC - positive control, AgPath - AgPath-ID™ One-Step RT-PCR kit (Thermo Fisher Scientific, Germany)

## Conclusions

To ensure a high level of diagnostic accuracy, it is highly recommended to pre-test each batch of PCR reagents thoroughly before applying it in routine diagnosis using more than 50 negative samples for specificity testing. Furthermore, it is of utmost importance to include also a reasonable number of appropriate controls such as NTCs, negative extraction controls and positive controls in every PCR run to minimize the risk of incorrect results further. Additional external quality assessment of the analytical results could be achieved by the participation in interlaboratory proficiency trials (FAO, 2015). Finally, in well-validated PCR-workflows, pooling of up to five samples might be an option for expanding capacities especially for the routine testing of low prevalences groups without any COVID-19-specific symptoms.

## Acknowledgments

We thank Bianka Hillmann, Katrin Schwabe and René Schöttner for excellent technical assistance. The study was supported by intramural funding of the German Federal Ministry of Food and Agriculture provided to the Friedrich-Loeffler-Institut.

## Ethical Statement

Anonymized human pharyngeal swab samples were obtained in the context of a COVID-19 monitoring study of the University Medicine of Greifswald (collaboration partner: Friedrich-Loeffler-Institut of Medical Microbiology): (SeCo study, registration number BB 068/20 by Ethics commission of the Greifswald University, Germany). The bovine oral or nasal swabs that were submitted to the Friedrich-Loeffler Institut for routine diagnostics were taken by the responsible farm veterinarians in the context of the health monitoring program of the respective farms, no permissions were necessary to collect the specimens. The unrelated cattle trial was reviewed by the responsible state ethics commission and was approved by the competent authority (permission number LALLF M-V/TSD/7221.3-2-016/17).

